# Methods in Cognitive Pupillometry: Design, Preprocessing, and Statistical Analysis

**DOI:** 10.1101/2022.02.23.481628

**Authors:** Sebastiaan Mathôt, Ana Vilotijević

## Abstract

Cognitive pupillometry is the measurement of pupil size to investigate cognitive processes such as attention, mental effort, working memory, and many others. Currently, there is no commonly agreed-upon methodology for conducting cognitive-pupillometry experiments, and approaches vary widely between research groups and even between different experiments from the same group. This lack of consensus makes it difficult to know which factors to consider when conducting a cognitive-pupillometry experiment. Here we provide a comprehensive, hands-on guide to methods in cognitive pupillometry, with a focus on trial-based experiments in which the measure of interest is the task-evoked pupil response to a stimulus. We cover all methodological aspects of cognitive pupillometry: experimental design; preprocessing of pupil-size data; and statistical techniques to deal with multiple comparisons when testing pupil-size data. In addition, we provide code and toolboxes (in Python) for preprocessing and statistical analysis, and we illustrate all aspects of the proposed workflow through an example experiment and example scripts.

---

The size of the eye’s pupil reflects a wide range of cognitive processes (reviewed in Beatty & Lucero-Wagoner, 2000; Loewenfeld, 1958; Mathôt, 2018). Increases in arousal (‘pupil-linked arousal’) or mental effort cause the pupil to dilate (i.e. become larger; Bradley et al., 2008; Hess & Polt, 1960; Kahneman & Beatty, 1966; Unsworth & Robison, 2014); and increases in brightness cause the pupil to constrict (i.e. become smaller) even if the source of brightness is merely imagined (Laeng & Sulutvedt, 2014), read about (Mathôt et al., 2017), covertly attended to (Binda et al., 2013a; Mathôt et al., 2013; Naber et al., 2013; Unsworth & Robison, 2017), or maintained in visual working memory (Husta et al., 2019; Zokaei et al., 2019). Pupil size is also often used as a non-invasive marker of activity of the locus coeruleus-norepinephrine (LC-NE) system (e.g. Aston-Jones & Cohen, 2005; de Gee et al., 2017; Joshi et al., 2016; Murphy et al., 2014). The measurement of pupil size to investigate cognitive processes is what we will refer to in this article as *cognitive pupillometry*.

There is no commonly agreed-upon workflow for conducting cognitive-pupillometry experiments: design criteria, preprocessing steps, and statistical analyses all differ vastly between studies. Some attempts at standardization have been made, but these are not easily applicable to cognitive pupillometry because they are too broad (Kelbsch et al., 2019), focused on a different subfield (e.g., listening effort; Winn et al., 2018), or focused on specific preprocessing steps (Kret & Sjak-Shie, 2018; Mathôt et al., 2018). Furthermore, guidelines are often conceptual rather than specific implementations, and the lack of concrete examples and code makes it difficult for researchers to incorporate these guidelines into their own workflow.

Here we aim to provide a hands-on, start-to-finish guide to cognitive pupillometry; that is, we provide concrete guidelines for designing cognitive-pupillometry experiments, for preprocessing of pupil-size data, and for conducting appropriate statistical analyses. All steps are illustrated with specific examples and code (in Python), which can be readily adapted to new experiments. In doing so, we will focus on what we consider to be a ‘typical’ cognitive-pupillometry experiment, which involves a computerized experiment with a trial-based structure in which the outcome of interest is a ‘task-evoked pupil response’ to a stimulus that is presented on each trial.

By limiting our focus to a ‘typical’ experiment, we can provide a comprehensive set of guidelines that cover every step of the process. But, of course, many experiments within cognitive pupillometry do not, or not exactly, follow this typical scenario. For example, to researchers interested in the relationship between sustained (tonic) pupil-size changes and tonic activity in the LC-NE system, pupil size during a pre-trial baseline may be a key measure of interest (e.g. Jepma & Nieuwenhuis, 2011; Pajkossy et al., 2017)—whereas for our purpose baseline pupil size is mainly a source of random noise; and to researchers interested in individual differences in cognitive abilities, resting-state pupil size as measured over a period of minutes may be a key measure of interest (e.g. Aminihajibashi et al., 2019; Tsukahara et al., 2016; Unsworth et al., 2021)—whereas for our purpose we are mainly interested in brief, task-evoked pupil responses. In other words, not all of our guidelines are directly applicable to all types of cognitive pupillometry; however, we believe that a thorough understanding of the basics (i.e. the topics covered in this manuscript) is useful in designing any kind of cognitive-pupillometry experiment.

Finally, our guidelines are meant to be illustrative, rather than prescriptive; that is, we show how things *could* be done—rather than how they *should* be done—in order to implement a workflow that is both easy to implement and that meets contemporary standards for good scientific practice. Our guidelines are intended to be a starting point for researchers who are interested in adopting (part of) our workflow, either by re-using our code directly or by implementing (part of) our workflow using different tools (e.g. Geller et al., 2020; Hershman et al., 2019; Kinley & Levy, 2021).

## Example experiment

Throughout this paper, we will refer to an experiment that we have recently conducted as an example. The theoretical motivation and methodological details for this experiment are described in Vilotijević and Mathôt (2022) and the associated preregistration (https://osf.io/ma4u9). Here, we will limit ourselves to those aspects of the experiment that generalize to many other experiments.

In brief, we asked whether pupil size increases as a function of attentional breadth; that is, we asked whether the pupil is larger when participants attend to the visual periphery as opposed to central vision. This question has been addressed several times before (Brocher et al., 2018; Daniels et al., 2012; Mathôt & Ivanov, 2019); however, previous studies were all limited by differences in task difficulty or visual input between conditions. Therefore, we set out to carefully re-investigate this question while controlling for all possible confounding variables.

Each trial started with the presentation of a symbolic cue that informed participants of where a target was likely to appear: in a central ring (near eccentricity), in a ring around this central ring (medium eccentricity), or in an outer ring (far eccentricity). The mapping between cue symbol and eccentricity (e.g. **□** → near, ○ → medium, ◁ → far) was randomly varied between participants. Next, the three rings appeared; each ring was filled with dynamic noise, that is, patches of oriented noise that changed every 30 Hz. At a random moment between 2000 and 3000 ms after the onset of the dynamic noise, a target (a subtle luminance increment or decrement) appeared, after which the dynamic noise continued for another 300 ms. On 80% of the trials, the target appeared at a random location within the cued ring; on 20% of the trials, the target appeared at a random location within either of the other two rings. Participants indicated whether they had seen a luminance increment or decrement by pressing the left or right key on the keyboard. Accuracy was maintained at 66% by a two-up-one-down staircase that manipulated the opacity of the target, separately for each eccentricity, using only responses from validly cued trials.

Our primary measure of interest was pupil size in the 3000 ms interval after the presentation of the cue and before the earliest possible presentation of the target. We predicted that pupil size should increase with increased cue eccentricity. Based on previous studies that investigated the effect of shifts of attention on pupil size, we expected this effect to emerge at the earliest 750 ms after the presentation of the cue (e.g. Mathôt et al., 2013); however, we did not have an a priori expectation about at which moment during the remaining 2250 ms interval the effect should emerge, nor for how long the effect should persist.

## Experimental design

A typical cognitive-pupillometry experiment follows the structure of a typical cognitive-psychology or cognitive-neuroscience experiment; that is, participants perform a computerized task that consists of many short trials during which stimuli are presented and responses are collected. By definition, one of the responses that is collected in a cognitive-pupillometry experiment is the pupil response; however, in many cases behavioral responses are collected as well. One or more independent variables are varied between trials. Our example experiment described above follows this structure. For a general introduction to experimental design for cognitive psychology and neuroscience, see Kingdom and Prins (2016). For the remainder of this article, we will assume that the reader is familiar with these kinds of experiments, and we will focus on those principles that are specific to cognitive-pupillometry experiments.

### Stimuli should ideally be constant between conditions

Light is the main determinant of pupil size (Mathôt, 2018); therefore, when using visual stimuli, brightness should be constant between conditions. (Assuming, of course, that stimulus brightness is not itself a manipulation of interest.) For example, when comparing pupil dilation in response to arousing and non-arousing images, the average brightness of the arousing and non-arousing images should be the same (e.g. Bradley et al., 2008). This is a minimum requirement for cognitive-pupillometry experiments, and is widely understood by researchers.

However, even when brightness is strictly controlled, other low-level differences between stimuli may still affect pupil size, and these are much more difficult to control. There are—and this is unfortunate for cognitive pupillometrists—very few stimulus properties that do *not* affect pupil size: any form of visual change or movement induces pupil constriction (Barbur et al., 1992; Mathôt, Melmi, et al., 2015; Slooter & van Norren, 1980; Ukai, 1985; Van de Kraats et al., 1977); color differences affect pupil size, generally such that blue stimuli result in a more sustained pupil constriction than red stimuli, though not necessarily in a more pronounced initial pupil constriction (Markwell et al., 2010); the distribution of brightness across the visual field affects pupil size, generally such that there is a bias towards foveal vision (Crawford, 1936; Hong et al., 2001), and to complicate matters further this foveal bias may be stronger for the initial phase of the pupil response than for more sustained pupil responses. And when it comes to auditory (Wang & Munoz, 2015), tactile (van Hooijdonk et al., 2019), and other sensory modalities, increased stimulus intensity generally results in increased pupil size.

Because of the many, hard-to-predict ways in which low-level stimulus properties affect pupil size, stimuli should ideally be kept strictly constant between conditions such that only the cognitive state of the participant is varied. This is sometimes referred to as the Hillyard principle, after Steven Hillyard, who advocated this principle for event-related-potential (ERP) experiments (see also Luck, 2005).

Our example experiment abides by the Hillyard principle: there is no difference in the dynamic noise between conditions; and because the mapping between cue symbol and cued eccentricity is randomly varied between participants, even slight potential differences in the pupil response to the different symbols, such as a slightly more pronounced pupil constriction in response to a triangle than to a circle, do not confound our results. (The location and size of the target do differ between conditions; however, the target only appears after the interval of interest.)

However, some research questions inherently require experiments that deviate from the Hillyard principle; for example, investigating the difference in the pupil response to arousing and non-arousing images requires comparing two different sets of images with each other. Therefore, in such cases the Hillyard principle cannot be upheld; however, it can still be approximated by matching visual stimuli between conditions on as many low-level properties as possible, such as brightness (at least), contrast, visual saliency, and the distribution of luminance across the display (for examples, see Binda et al., 2013b; Naber & Nakayama, 2013). There are no standardized tools to do this. However, a straightforward way to match visual stimuli on brightness and contrast is to equate the average pixel value (as a measure of brightness) and the standard deviation of pixel values (as a measure of contrast). An example script is provided at https://osf.io/ngq5b/.

Importantly, matching is always approximative and never perfect. Therefore, a key criterion is that a reasonable and informed person, such as a good peer reviewer, should agree that, while some confounds likely remain, they are unlikely to affect the results.

### Eye position should ideally be constant between conditions

The position and movement of the eyes affect pupil size in two main ways. First, for most eye trackers, the angle from which the camera records the eye affects the recorded (as opposed to the actual) size of the pupil. This eye-position-based distortion of pupil size is often called the pupil-foreshortening error (PFE), and directly results from the fact that the surface of the pupil appears smaller in the camera image when it is recorded from the side, and thus looks like an ellipse, as compared to when it is recorded from the front, and thus looks like a circle. Some eye trackers are more susceptible to the PFE than others (Petersch & Dierkes, 2021), and there are ways to compensate for the PFE algorithmically during data analysis (Brisson et al., 2013; Gagl et al., 2011; Hayes & Petrov, 2016); however, although the PFE can certainly be minimized, you can never assume that pupil-size recordings are completely unaffected by the angle from which the eye tracker records the pupil.

Second, pupil size may really be affected by the position of the eye, independent of the PFE. This may happen when, say, the participant is seated such that it is more comfortable to look at the lower part of the display than at the upper part of the display (Mathôt, Melmi, et al., 2015).

The increased effort or agitation associated with looking up may then result in pupil dilation. This is distinct from the PFE in the sense that it is not a recording artifact but a legitimate change in pupil size; however, even though it is real, it may still be a confounding factor in experiments in which eye position is not controlled.

Third, eye movements are followed by pupil constriction (Knapen et al., 2016; Mathôt, Melmi, et al., 2015; Zuber et al., 1966), presumably as a result of the changes in visual input that are caused by eye movements. Constriction also occurs after eye blinks, which are similarly accompanied by a sudden change in visual input due to the closing and opening of the eyelid. This post-eye-movement and post-blink constriction can be compensated for algorithmically during data analysis (Knapen et al., 2016; Yoo et al., 2021); however, just as for the PFE, you can never assume that pupil size is completely unaffected by eye movements and blinks.

Because of these considerations, eye position should ideally be constant between conditions. In our example experiment, we ensured this by asking participants to keep their eyes fixated on a central stimulus throughout the experiment, thus minimizing eye movements. In some experiments in which it is necessary for participants to make eye movements, eye position can be strictly matched between conditions (Mathôt, Melmi, et al., 2015). However, in other experiments, for example experiments involving reading or other forms of natural eye movements, it is not feasible to strictly match eye position between conditions. In that case, eye position should be matched as closely as possible (Gagl et al., 2011). A key criterion is once again that a reasonable and informed person should agree that eye position is unlikely to affect the results.

### Trials should ideally be slow-paced

The pupil light response (PLR) is the fastest pupil response with a latency of about 200 ms; that is, the pupil starts to constrict around 200 ms after stimulus onset, and reaches maximal constriction around 750 ms after stimulus onset. Therefore, experimental manipulations that affect the strength and/or latency of the pupil light response to a stimulus tend to arise within 500 ms (Binda & Murray, 2015; Olmos-Solis et al., 2018; Wilschut & Mathot, 2022). Another kind of pupil response that emerges rapidly is the orienting response that occurs when something captures attention; the orienting response is accompanied by a pupil dilation that peaks around 500 ms (Mathôt et al., 2014; Wang & Munoz, 2014). Therefore, the oft-heard claim that pupil responses are very slow is not *necessarily* true.

However, what is true is that most cognitive-pupillometry experiments focus on pupil responses that do arise slowly; specifically, most experiments focus on pupil dilation as a marker of arousal or effort (e.g., Bradley et al., 2008; Rieger et al., 2015; Urai et al., 2017; Zekveld et al., 2010). If there is a clearly defined triggering stimulus, such as a burst of noise, pupil size tends to peak around 1 s after stimulus onset (Hoeks & Levelt, 1993). However, in many studies, the trigger is an internal cognitive process without a clearly defined temporal onset, such as a challenging calculation (Hess & Polt, 1964; Stoll et al., 2013) or a spoken sentence that is hard to understand (Koelewijn et al., 2012). In those cases, the resulting pattern of pupil dilation is hard to predict, but tends to be even slower and more diffuse.

Because of these considerations, cognitive-pupillometry experiments should ideally be slow-paced. However, experiments should not be *too* slow-paced for the simple reason that, as a researcher, you also need to be able to collect a sufficient amount of data within a reasonable time. As a rule of thumb, we suggest having the stimulus of interest followed by an interval of 2 - 3 s during which no other events happen (see also below), and which serves as the period of interest for pupil-size analysis. We suggest an intertrial interval of at least 3 s, to reduce (but not eliminate!) carry-over effects in the pupil response from one trial to the next.

For this reason, in our example experiment, there was at least 3000 ms between the onset of the cue and the onset of the target stimulus. The intertrial interval, as measured from the response on one trial until the onset of the cue on the next trial, was around 4 - 6 s; this is longer than necessary for the purpose of pupillometry, but resulted from the fact that stimulus generation occurred during this interval and took a substantial amount of time.

### Pupil size should be ideally be measured while participants do nothing

An increase in physical effort is accompanied by pupil dilation. This is true for intense physical exercise, and to a lesser-yet-measurable extent also for pressing a key on a keyboard, making a saccadic eye movements towards a target, pressing a button on a gamepad—or more generally any action through which participants respond in an experiment. Combined with the fact that response times (RTs) often vary between conditions, this response-locked pupil dilation can be a confounding factor when not properly accounted for.

To illustrate this issue, consider a hypothetical experiment in which participants indicate whether a character string corresponds to a word or not (i.e. a lexical-decision task); the difficulty of the word is the independent variable (difficult vs easy) and the pupil response to the word is the dependent variable. You might reasonably predict that processing a difficult word takes more mental effort, and should thus result in stronger pupil dilation, as compared to an easy word. (And given the proper experimental design this is quite likely what you would find.) However, participants also respond more slowly to difficult words than to easy words. Because of this difference in RT, you will likely find that the pupil initially dilates more strongly in response to easy as compared to difficult words—simply because of the response-locked pupil dilation.

Because of this, a task-evoked pupil response should not be recorded while participants are also giving a response. Rather, the response should be delayed until sometime after. In a lexical-decision experiment, this could mean that the character string is followed by a period of about 2 s of passive viewing during which pupil size is recorded; only after this period do participants indicate whether they saw a word or a non-word (for similar approaches, see Einhäuser et al., 2010; Snell et al., 2018). In our example experiment, we avoided response-locked pupil dilation by measuring pupil size during the interval preceding the target, before participants could even prepare a response, thus avoiding all motor preparation.

### Ambient lighting should ideally be intermediate and matched to display brightness

There are four main concepts that are all related to light but each reflect a different aspect. The total amount of light that is emitted by a light source, or *luminous flux*, is expressed in lumen (or candela); this measure is not often reported in experimental settings, because it does not correspond to something that is directly observable (i.e. you cannot observe the light that a lamp emits in the direction away from you).

The amount of light that falls from any direction onto a surface (such as the pupil), or *illuminance*, is expressed in lux; this measure is often reported in experimental settings as an indication of the ambient light level in a laboratory, and is the combined result of ceiling lights, lamps, the background luminance of the display, and other sources of light in the laboratory. Illuminance can be measured with an illuminance meter, which is generally a small hand-held device. To measure illuminance as it would be from the perspective of the participant, place the light sensor where the participant’s head would be and orient it towards the screen. For our example experiment, illuminance was 33 lux.

The amount of light that is emitted by a light source (such as a visual stimulus) in a specific direction (such as towards the pupil), or *luminance*, is expressed in candela per square meter (cd/m^2^); this measure is often reported in experimental settings as an indication of the intensity of a visual stimulus. Luminance can be measured with a luminance meter (or photometer), which is also a small handheld device, but is distinct from the illuminance meter; that is, you can generally not measure illuminance and luminance with the same device. To measure the luminance of a visual stimulus, first present the stimulus on the screen; next, point the sensor towards the stimulus from a close distance or (depending on the model) place the sensor on the screen while covering the stimulus with the device. For our example experiment, the luminance of both the background and the stream of dynamic noise was 33.1 cd/m^2^.

Finally, brightness refers to the subjective impression of luminance, and as such does not have a natural unit. Although luminance and brightness strictly speaking refer to different concepts, many authors, including ourselves on many occasions (such as in the heading above), use the terms interchangeably.

Pupil size varies between roughly 2 and 8 mm in diameter (Mathôt, 2018; Pan et al., 2022), depending mainly on the amount of light that enters the eye. Both illumination and luminance play a role: roughly speaking, the illumination of the laboratory determines baseline pupil size, whereas the luminance of the stimuli determines the strength of visually evoked pupil responses (see Stimuli should ideally be constant between conditions).

For most cognitive-pupillometry experiments, pupil size should not approach the physiological lower and upper limits of 2 and 8 mm, for the simple reason that maximally constricted or dilated pupils cannot constrict or dilate any further in response to experimental manipulations. Experiments should therefore not be conducted in a very dark or very bright environment (unless there are specific reasons to do so, for example if the experiment requires dark-adapted pupils). However, within a broad range of baseline pupil sizes, and thus for a broad range of illuminance levels, task-evoked pupil responses are reliably observed (Bradshaw, 1969; Cherng et al., 2020; Pan et al., 2022; Peysakhovich et al., 2017; Reilly et al., 2019; Steinhauer et al., 2004). As a rule of thumb, baseline pupil sizes between 3 and 7 mm are suitable. In our dimly illuminated laboratory of 33 lux, baseline pupil sizes were around 3.5 mm. Some eye trackers only provide pupil size in arbitrary units without an easy way to convert these units to millimeters of diameter (see If necessary: converting pupil size from arbitrary units to millimeters of diameter); in this case, even without knowing the exact pupil size of the participants, you can safely aim for an illumination level that is dim but still allows you to read comfortably.

It is also important to use a display luminance that roughly matches the illuminance of the room; more specifically, using a bright stimulus display in an otherwise dark room results in discomfort glare, which is the unpleasant sensation that results from a small yet intense light source. Discomfort glare is accompanied by a pronounced decrease in pupil size (Tyukhova & Waters, 2019). The intermediate luminance (33.1 cd/m^2^) of the stimulus display in our example experiment looked subjectively natural given the illumination of the room.

In sum, ambient lighting should ideally be intermediate; fortunately, because a broad range of illuminance levels is acceptable, no special equipment is needed to find the ‘correct’ illuminance. Similarly, display brightness should ideally match the ambient lighting, but since the main criterion is that the display does not subjectively stand out from its environment, no special equipment is again needed to find the ‘correct’ luminance. However, it is convenient to nevertheless have illuminance and luminance (or photo-)meters, because these allow you to measure and report the exact conditions under which your experiment took place.

### All data should ideally be stored in a single file per participant

A typical cognitive-pupillometry experiment is implemented using experiment-building software that is connected to an eye tracker that records gaze position and pupil size. In our example experiment, we used OpenSesame (Mathôt et al., 2012) to implement the experiment, PyGaze (Dalmaijer et al., 2014) to implement eye tracking in the experiment, and an EyeLink eye tracker to record pupil size. In set-ups of this kind, there are two distinct data files: the data file of the experiment-building software, in our case the OpenSesame `.csv` file, which contains experimental variables, etc.; and the data file of the eye tracker, in our case the EyeLink `.edf` file, which contains eye-movement and pupil-size data. It is possible to cross-reference these two data files afterwards during data analysis; however, doing so is cumbersome and error-prone.

Therefore, it is good practice to log all data to a single file already during the experiment; usually, this single file is the eye-tracker data file, because it is more convenient to send time stamps, experimental variables, and other relevant information from the experiment-building software to the eye tracker, than it is to send all eye-position and pupil-size data from the eye tracker to the experiment-building software. In our example experiment, we did this by marking the start and end of relevant intervals (cue interval, dynamic-noise interval, etc.) in the EyeLink `.edf` file, and by logging all experimental variables (cue eccentricity, participant response, etc.) to the EyeLink `.edf` file at the end of each trial.

Finally, it is important to decide in advance how you intend to analyze the data (what analysis software you will use, what kind of information you need to extract, etc.), and to log the data in a format that makes this as easy as possible. In our example experiment, we used the Python EyeLinkParser module^1^; this module automatically processes variables that have been logged with `var [name] [value]` messages in the `.edf` file, and we therefore designed the experiment to send messages in this format. In contrast, had we intended to analyze the data with the EyeLink DataViewer software, we would instead have sent `!V TRIAL_VAR [name] [value]` messages, which is how DataViewer assumes that variables are logged.

## Preprocessing

Here we use the term ‘preprocessing’ to refer to all steps involved in transforming the raw data, as it is stored during the experiment, into a format that is suitable for statistical analysis and visualization. The order matters, and the preprocessing steps below are described in the order in which they should be performed.

### Parsing raw data into an analysis-friendly data structure

Preprocessing starts with a set of raw data files, one for each participant, that contain all relevant data; this data includes experimental variables, start and end markers for trials and relevant intervals, and gaze-position and pupil-size data. In our example experiment, this data file is the EyeLink `.edf` file, but every brand of eye tracker uses its own custom format.

The first step is to parse the raw data into a data structure that is suitable for further processing. This generally involves a programming library or user interface that has been specifically designed for this purpose. For example, GazeR is an R library that parses raw eye-tracking data into an R `data.frame` object for further analysis (Geller et al., 2020). CHAP (Hershman et al., 2019) and PuPl (Kinley & Levy, 2021) are MATLAB-based user interfaces that parse raw data into custom data structures that are used internally for further processing.

For our example experiment, we used the Python EyeLinkParser library to parse the raw data into a Python `DataMatrix` object^2^. Assuming that data has been logged following the assumptions of EyeLinkParser (see All data should ideally be stored in a single data file), the parser automatically recognizes experimental variables, and splits eye-position and pupil-size into separate intervals or ‘epochs’ (e.g. cue interval, dynamic-noise interval, etc.). This results in a spreadsheet-like data structure in which each row corresponds to a single trial, and each column corresponds to a single variable (Fig. 3). For variables that contain a single value per trial, such as `response_time`, one cell in the `DataMatrix` contains a single value. However, for variables that contain time series, such as pupil size during the cue interval (`pupil_cue`), one cell contains a series of values. You can think of this as a spreadsheet in which certain columns, those containing time series (`SeriesColumn` objects), stick out of the spreadsheet to accommodate their additional dimension: time.

**Figure 1.**
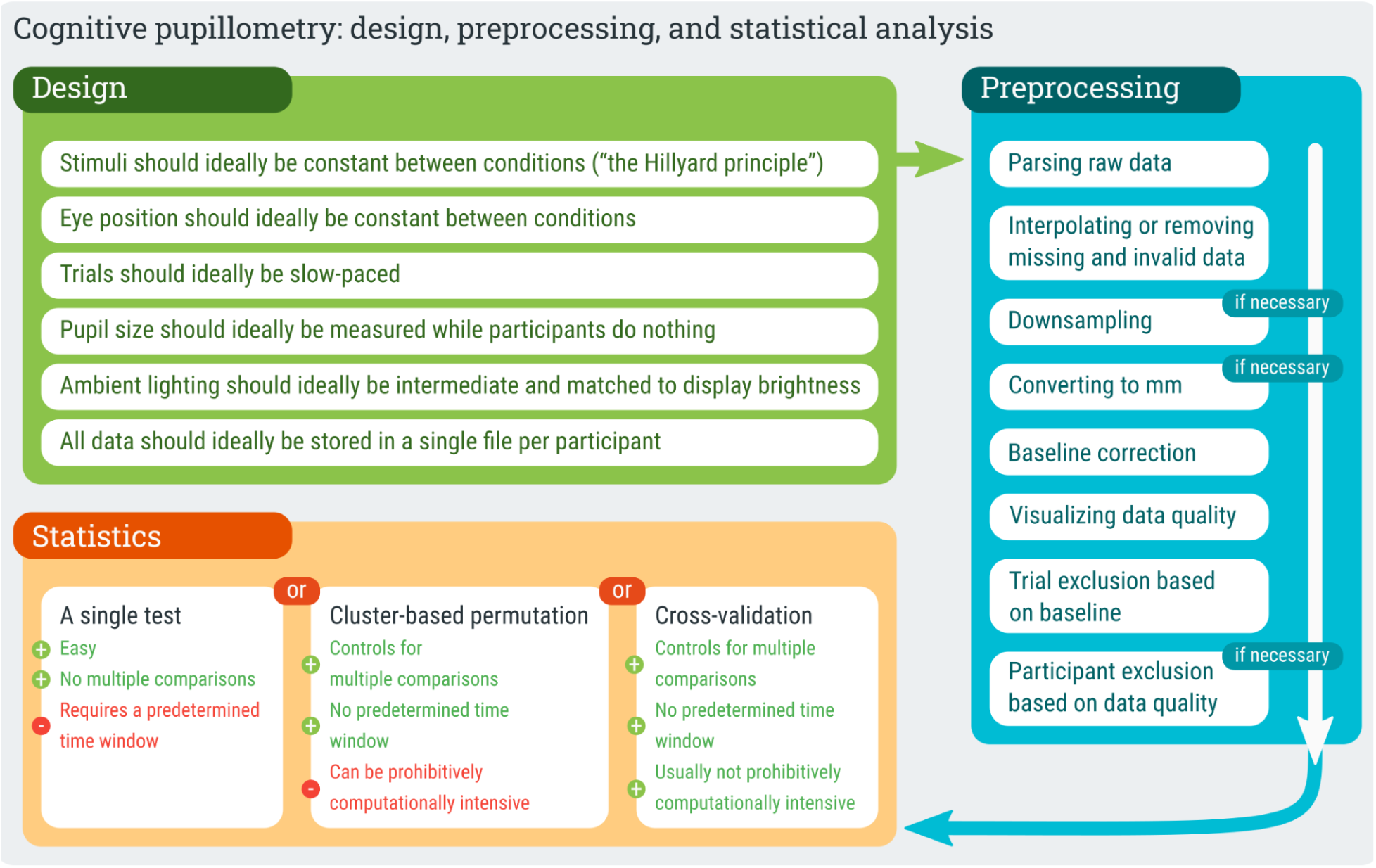
A schematic representation of the proposed workflow for conducting cognitive-pupillometry experiments.

**Figure 2.**
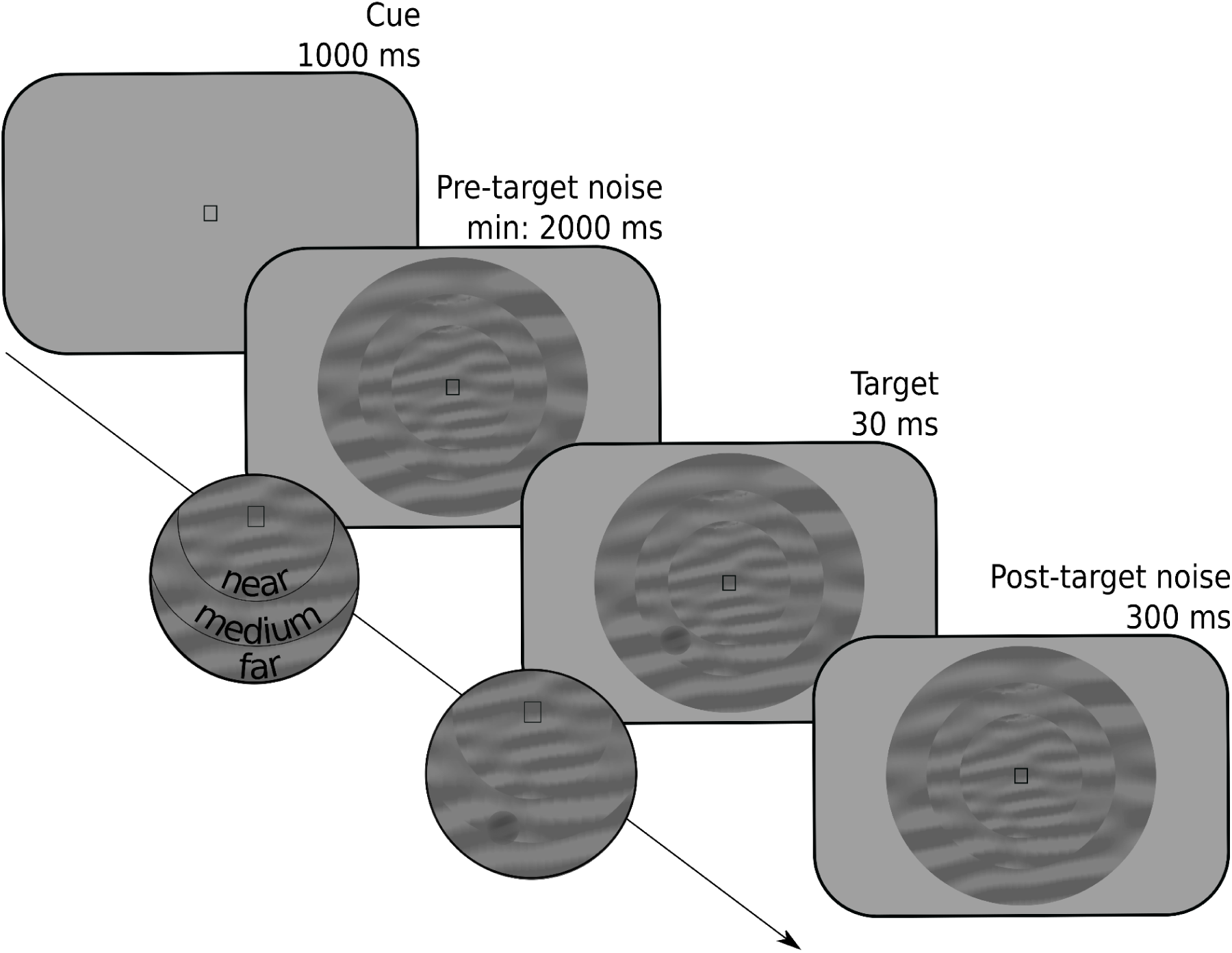
Schematic trial sequence for the example experiment. Participants reported whether a target was a luminance increment or, as in this example, a luminance decrement embedded in a dynamic stream of noise. The probable eccentricity (near, medium, far) of the target was indicated by a symbolic cue (a square in this example).

**Figure 3.**
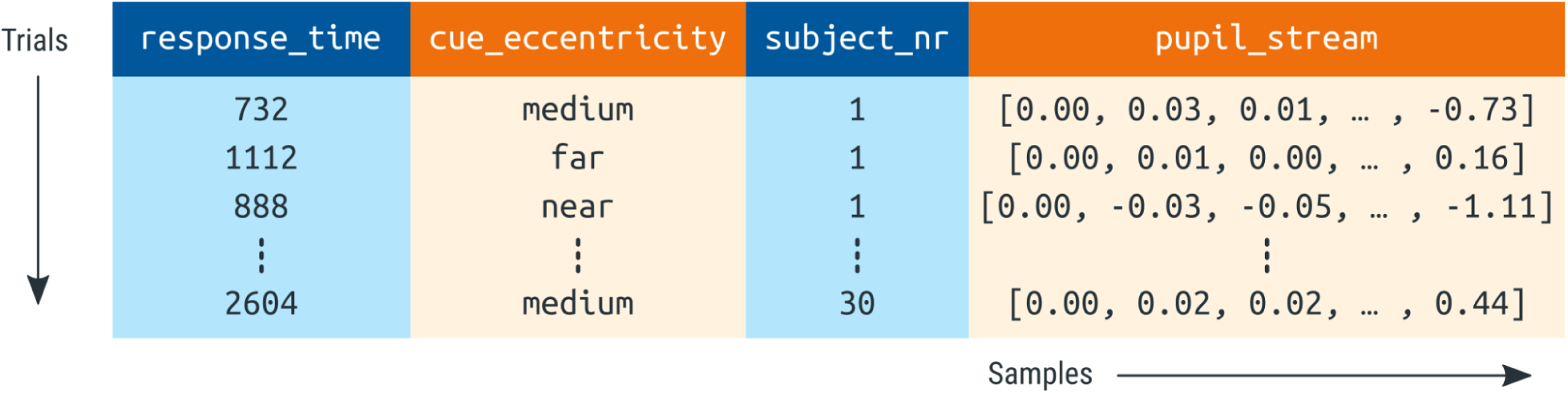
A schematic representation of a `DataMatrix` object. Each row represents a trial. Each column represents a variable. Pupil-size data (here: `pupil_stream`) is stored in a special column type (`SeriesColumn`) that has an additional dimension to store how pupil size changes as a function of time (sample number).

The fact that time-series data (e.g. pupil size) can co-exist with single-value data (e.g. response time) in a single `DataMatrix` object is convenient for further analysis. However, this is only one of many ways in which time series can be represented. Another common way is using a two-dimensional spreadsheet in which each row corresponds to one sample, while other trial-level variables, such as response time, are duplicated for all samples from the same trial. (Analogously to how participant-level variables, such as subject number, are duplicated for all trials of the same participant in Fig. 3.) The main advantage of this kind of representation is that it is compatible with commonly used data structures, such as `data.frame` in R and `pandas.DataFrame` in Python. The main disadvantages are that this representation consumes much more memory than necessary (due to massive data duplication) and makes it less straightforward to perform certain common operations, such as averaging a time series across trials.

### Interpolating or removing missing and invalid data (e.g. due to blinks)

Missing data occurs when the eye tracker fails to record any pupil size at all. This can happen when the participant moves slightly, causing the eye tracker to lose the eye, when there is a technical malfunction of the eye tracker, or when the participant has fully closed the eyes. Most eye trackers indicate missing data with a special value; in the case of the EyeLink, missing data is initially indicated with the value 0. During parsing, missing data should ideally be recoded to `NaN` values (not a number), which is a standard way to represent missing or invalid data.

Invalid data occurs when the eye tracker does record pupil size, but the measured pupil size does not accurately reflect the actual pupil size. This can happen when the participant is in the process of blinking, such that the pupil is partly obscured by the eyelid, or when the eye tracker does not reliably extract the entire pupil from the camera image. Unlike missing data, invalid data is not represented in any special format, and can only be detected indirectly, through various quality-check measures.

Pupil-size data, even when it is of high quality, invariably contains missing and invalid data (discussed also in Kret & Sjak-Shie, 2018; Mathôt et al., 2018). Therefore, an important step in preprocessing is to first identify invalid data and remove it, that is, to turn invalid data into missing data. In principle, this is sufficient, because missing data does not cause measurement error in the way that invalid data does. However, missing data does reduce statistical power, and can make visualizations of average pupil data look less smooth. For these reasons, it is good practice to interpolate missing data whenever this is reasonably possible.

The goal of interpolation is to estimate missing or invalid data by drawing a line through valid data points. This is usually done using either a linear interpolation, which draws a straight line between two valid data points, or using quadratic (cubic-spline) interpolation, which draws a smooth line through four valid data points, which results in interpolations that more closely resemble natural pupil-size changes. There are various ways in which interpolation of missing pupil-size data can be implemented; such procedures are sometimes referred to as ‘blink reconstruction’, since blinks are the primary reason for missing data (for alternative implementations, see Hershman et al., 2019; Kinley & Levy, 2021; Kret & Sjak-Shie, 2018). Here we will focus on the ‘advanced’ algorithm as implemented in the DataMatrix `blinkreconstruct()` function. This algorithm is based on the procedure described in Mathôt (2013), and has been optimized to catch many of the edge cases in which the original algorithm failed.

The algorithm uses a recursive procedure in which it first reconstructs pupil size during blinks. To do so, it identifies the onset and offset of a blink based on a velocity threshold; that is, a blink is assumed to start when pupil size rapidly decreases (due to closing of the eyelid, thus rapidly obscuring the pupil), and to end when pupil size stabilizes again after a period of rapid pupil-size increase (due to opening of the eyelid). A blink is generally preceded and followed by several milliseconds of unreliable data; therefore, a margin (by default 10 ms) around the blink is also marked as missing. A blink cannot be longer than a certain duration (by default 500 ms); longer blink-like periods are not reconstructed, based on the intuition that interpolation only makes sense for brief periods during which pupil size is predictable.

Once the onset and offset of a blink have been determined, the algorithm first tries to perform cubic-spline interpolation; this requires two additional points, equally spaced before the onset of the blink and after the offset of the blink. If these points can be determined, missing data during the blink is interpolated with a smooth curve (Fig. 4a). If these points cannot be determined, for example because one of the points falls outside the time series or inside another blink, the blink is interpolated linearly with a straight line (Fig. 4b).

**Figure 4.**
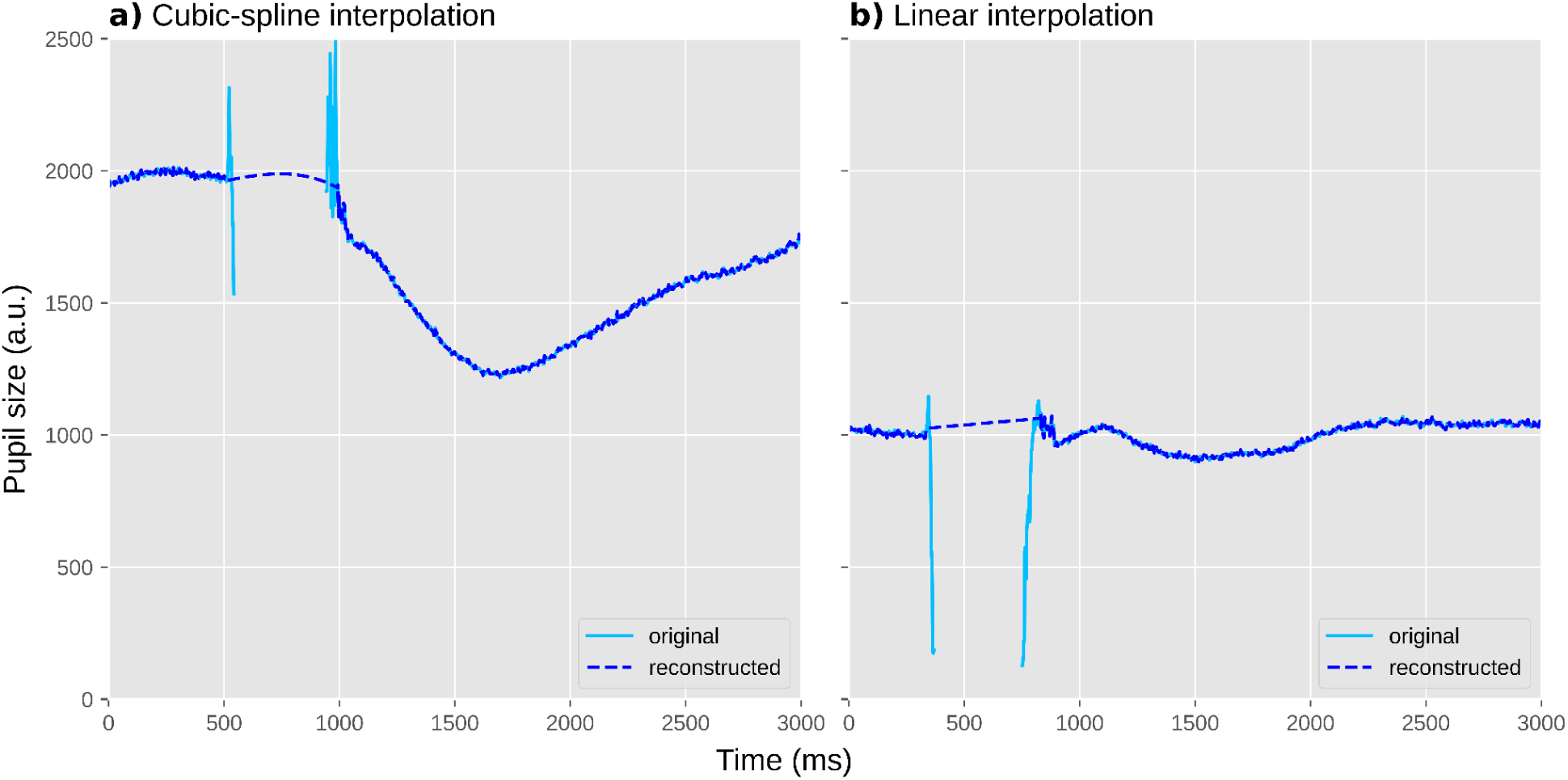
Two individual trials during which a blink occurred. Blinks are often characterized by a sharp drop in pupil size (due to closing of the eyelid) followed by a period of missing data (due to the eyelid being fully closed), followed in turn by sharp increase in pupil size (due to the eyelid re-opening). a) An example of cubic-spline interpolation, based on four points around the blink. This results in a smooth curve. b) An example of linear interpolation, based on two points around the blink. This results in a straight line.

Once a blink has been identified and reconstructed, the algorithm starts again from scratch (i.e. it is a recursive procedure) until no blinks are identified anymore. At this point, any remaining invalid data is marked as missing. This is done by removing all pupil-size data points that deviate more than a threshold from the mean pupil size of the series (by default ±3SD) or where pupil size (very) rapidly increases or decreases. Finally, margins (by default 20 ms) around all periods of missing data are similarly marked as missing.

### If necessary: downsampling

The latency of the pupil light response is around 200 ms, and the earliest cognitive effects on pupil size emerge around 500 ms after the triggering stimulus (see Trials should ideally be slow-paced). This means that a sampling rate of 20 Hz, such that pupil size is measured once every 50 ms, is sufficient when, as in most cognitive-pupillometry experiments, you are interested in how pupil size differs between experimental conditions. (A higher sampling rate is desirable when you are interested in measures such as maximum constriction velocity, which requires many measurements during the fast initial constriction of the pupil light response.) For visualization purposes, a higher sampling rate is desirable to be able to generate smooth figures of pupil size over time; however, even for visualization, a sampling rate of more than 100 Hz is rarely useful.

Therefore, if you are using an eye tracker that records at a higher sampling rate, it is convenient to downsample the signal to 100 Hz. The main reason for doing so is to reduce memory consumption, which without downsampling can easily become prohibitive for large datasets. In the case of our example experiment, we downsampled the original 1000 Hz recording by a factor of 10 to 100 Hz. Importantly, downsampling should be performed *after* preprocessing steps that benefit from a high sampling rate, such as blink reconstruction.

### If necessary: converting pupil size from arbitrary units to millimeters of diameter

For most cognitive-pupillometry experiments, the measure of interest is not the size of the pupil in absolute terms, but rather the change in pupil size between conditions. As such, the unit in which pupil size is reported is not crucially important. Nevertheless, many researchers prefer to express pupil size in millimeters of diameter. Doing so makes it possible to verify that pupil size does not approach physiological limits (see Ambient lighting should ideally be intermediate and matched to display brightness), and makes it possible to compare effect sizes in natural units: a pupil constriction of 4 mm in response to a flash of light is in an entirely different class of effect sizes than a pupil dilation of 0.1 mm in response to a challenging calculation, even though both may be highly significant when tested statistically.

Some eye trackers, such as the Tobii series of eye trackers, automatically report pupil size in millimeters; they are able to do so because the eye tracker contains a depth sensor that estimates the distance between the eye tracker and the participant’s face, which in turn allows the eye-tracking software to transform pupil size from camera-image-specific units (e.g. pixel count, or the size of the longest axis of the best fitting ellipse in pixels) to millimeters. However, many eye trackers, such as the EyeLink, do not offer this functionality and simply express pupil size in arbitrary units. In that case, it may be possible to determine a conversion formula yourself.

In the case of our example experiment, our group had previously determined a conversion formula for our laboratory set-up, allowing us to convert the EyeLink’s arbitrary units to millimeters (Wilschut & Mathot, 2022). To do so, we printed fifteen black circles of various known sizes on a white sheet of paper and held this in front of the eye tracker just above the chinrest. The EyeLink accepts these circles as pupils, and reports a size for them. This provided us with ground-truth pupil sizes (i.e. the size of the circles as measured with a ruler) and recorded pupil sizes in arbitrary units, from which we derived a conversion formula with the help of an online application (such as https://mycurvefit.com) that automatically determines the best-fitting function given a set of predictors and observations.

### Baseline correction

Pupil size fluctuates in waves of several seconds that reflect the waxing and waning of arousal (Mathôt, Siebold, et al., 2015; Reimer et al., 2014), and in slower waves that reflect more prolonged states of fatigue and wakefulness (Geacintov & Peavler, 1974; Lowenstein et al., 1963). In cognitive-pupillometry experiments that are not primarily concerned with general states of arousal and wakefulness, such fluctuations are a source of noise that reduce statistical power to detect differences in task-evoked pupil responses between conditions.

Baseline correction is a technique to remove the impact of trial-to-trial fluctuations in pupil size (Mathôt et al., 2018). This is done for each trial separately, usually by subtracting the mean pupil size during a ‘baseline period’ from all subsequent pupil-size measurements (subtractive baseline correction; recommended), or sometimes by dividing all subsequent pupil-size measurements by the mean baseline pupil size (divisive baseline correction; not recommended, see Mathôt et al., 2018). As a result, pupil size starts from 0 (for subtractive correction) or 1 (for divisive correction) during the baseline period on every trial, and only the change in pupil size—the task-evoked pupil response—remains.

Crucially, a baseline period should itself not be affected by the experimental manipulations. In practice, this means that the baseline period should come before the stimulus that triggers the task-evoked pupil response; alternatively, the baseline period can coincide with the onset of the triggering stimulus, as long as the duration of the baseline period is below the minimum latency of the pupil response (i.e. less than 200 ms). Also, the baseline period should not contain too many blinks or other artifacts that could affect baseline pupil size (see Excluding trials based on baseline pupil size); therefore, we prefer to use a short baseline period, since this reduces the chance of artifacts. In our example experiment, we used the first 50 ms after the onset of the cue as the baseline period, because the cue is the stimulus that triggers the shift in attentional breadth that we hypothesized would affect pupil size.

A subtle point related to baseline correction that is often overlooked (including by us, and we thank an anonymous reviewer for bringing this to our attention) is that it introduces counter-intuitive contingencies between pupil size *during* the baseline period and baseline-corrected task-evoked pupil responses *after* the baseline period; these contingencies at least, but not necessarily exclusively, take the form of regression to the mean. Specifically, when pupils are large during the baseline period, they are likely to decrease in size afterwards, simply because by chance large pupils are more likely to become smaller than they are to become even larger. Conversely and for the same reason, when pupils are small during the baseline period, they are likely to increase in size afterwards. This means that, simply as a result of regression to the mean, there is likely to be a negative correlation between pupil size during the baseline and the strength of the subsequent baseline-corrected task-evoked pupil response, as has in fact been reported in several studies (e.g. de Gee et al., 2014; Gilzenrat et al., 2010). Because of such contingencies, researchers should be careful when interpreting correlations between baseline pupil size and baseline-corrected task-evoked pupil responses, and ensure that such correlations—which may be theoretically important—are not statistical artifacts (see Mridha et al., 2021 for an example of correcting for regression to the mean).

### Verifying and visualizing data quality

In an ideal world, pupil-size data consists of smooth traces that are unaffected by blinks, eye movements, or recording artifacts. In addition, in this ideal world, gaze position is perfectly constant while pupil size is being recorded. However, in the *real* world, the quality of pupil-size data never attains this ideal. Therefore, it is important to get a sense of the quality of your dataset by visualizing relevant aspects of the data.

A useful visualization is to plot all pupil-size traces as semi-transparent lines in a single figure (Fig. 7a); good-quality data will look like a tangle of lines. Another useful visualization is a histogram of baseline pupil sizes (Fig. 7b); good-quality data will be roughly normally distributed, and often has a slight skew that can either be to the left (as in Fig. 7b; less common) or to the right (more common). These figures are best created separately for each participant to avoid clutter, and collapsed across experimental conditions to avoid yourself from being biased by whether the data show the desired effect when assessing data quality.

**Figure 5.**
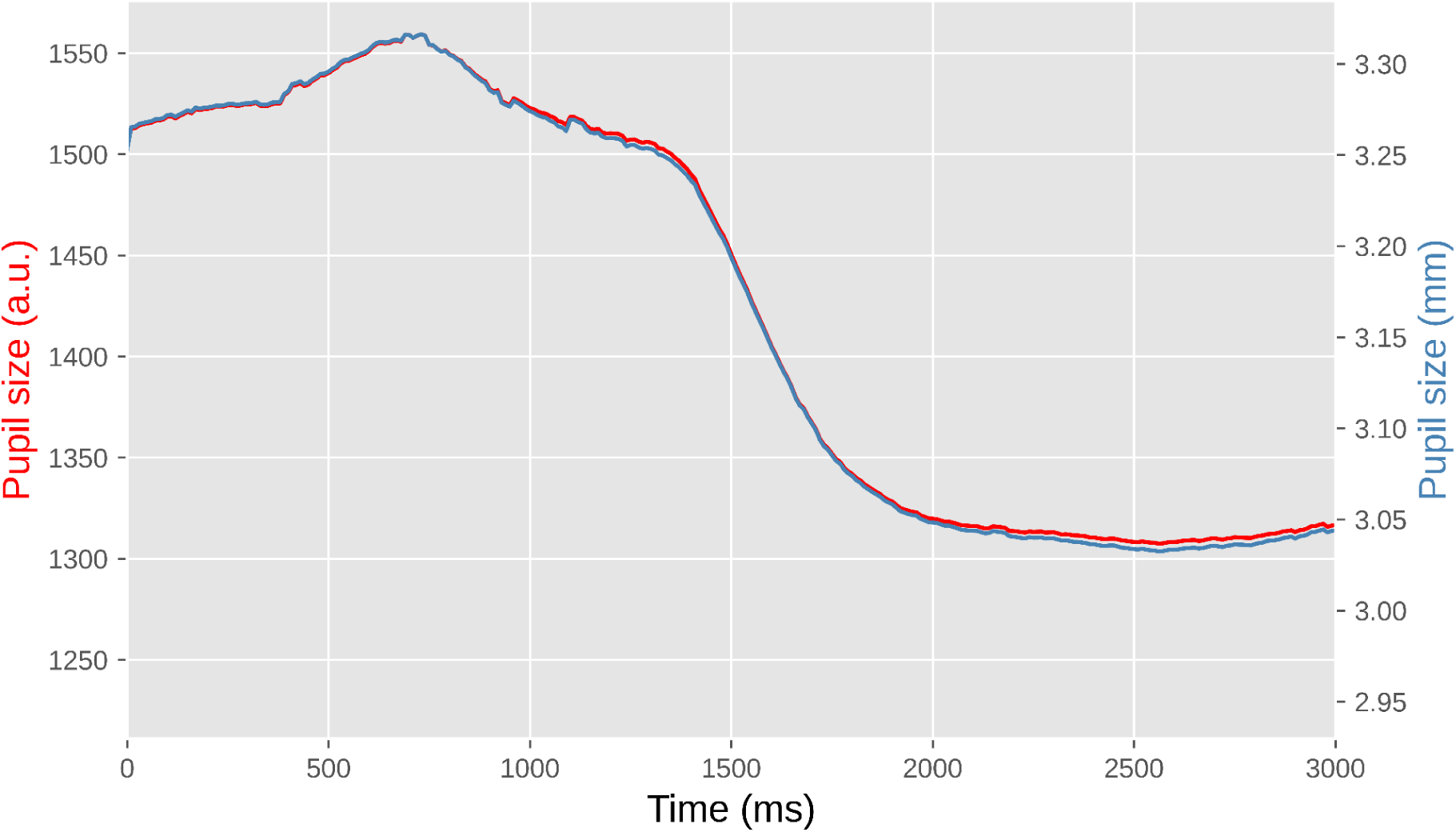
Pupil size during a single trial from our example experiment. The left y-axis shows pupil size in arbitrary units as recorded by the EyeLink eye tracker. The right y-axis shows pupil size in millimeters of diameter based on a conversion formula. The lines do not overlap perfectly because the conversion is not linear.

**Figure 6.**
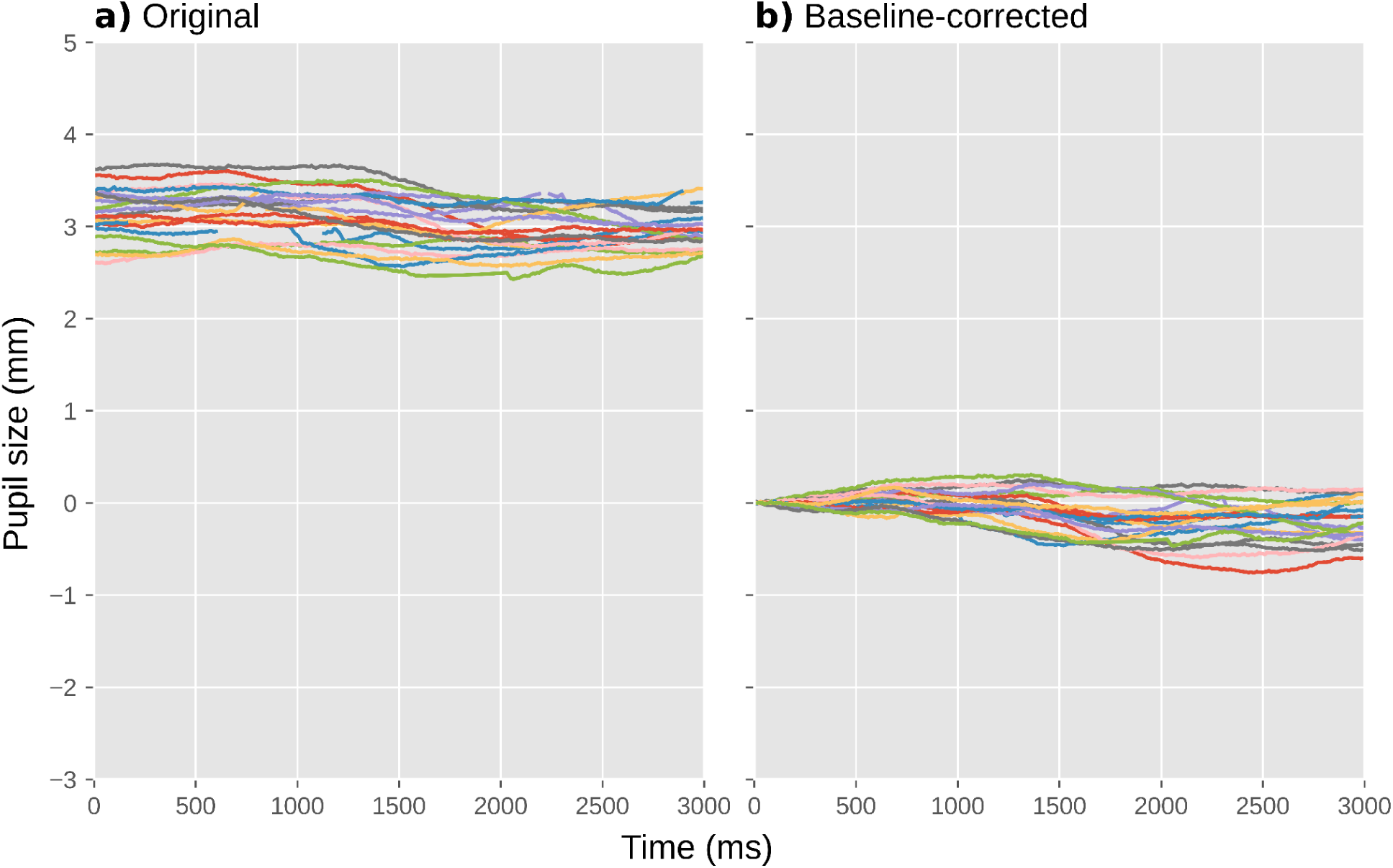
Pupil size during a selection of individual trials from our example experiment. a) Before baseline correction. b) After baseline correction.

**Figure 7.**
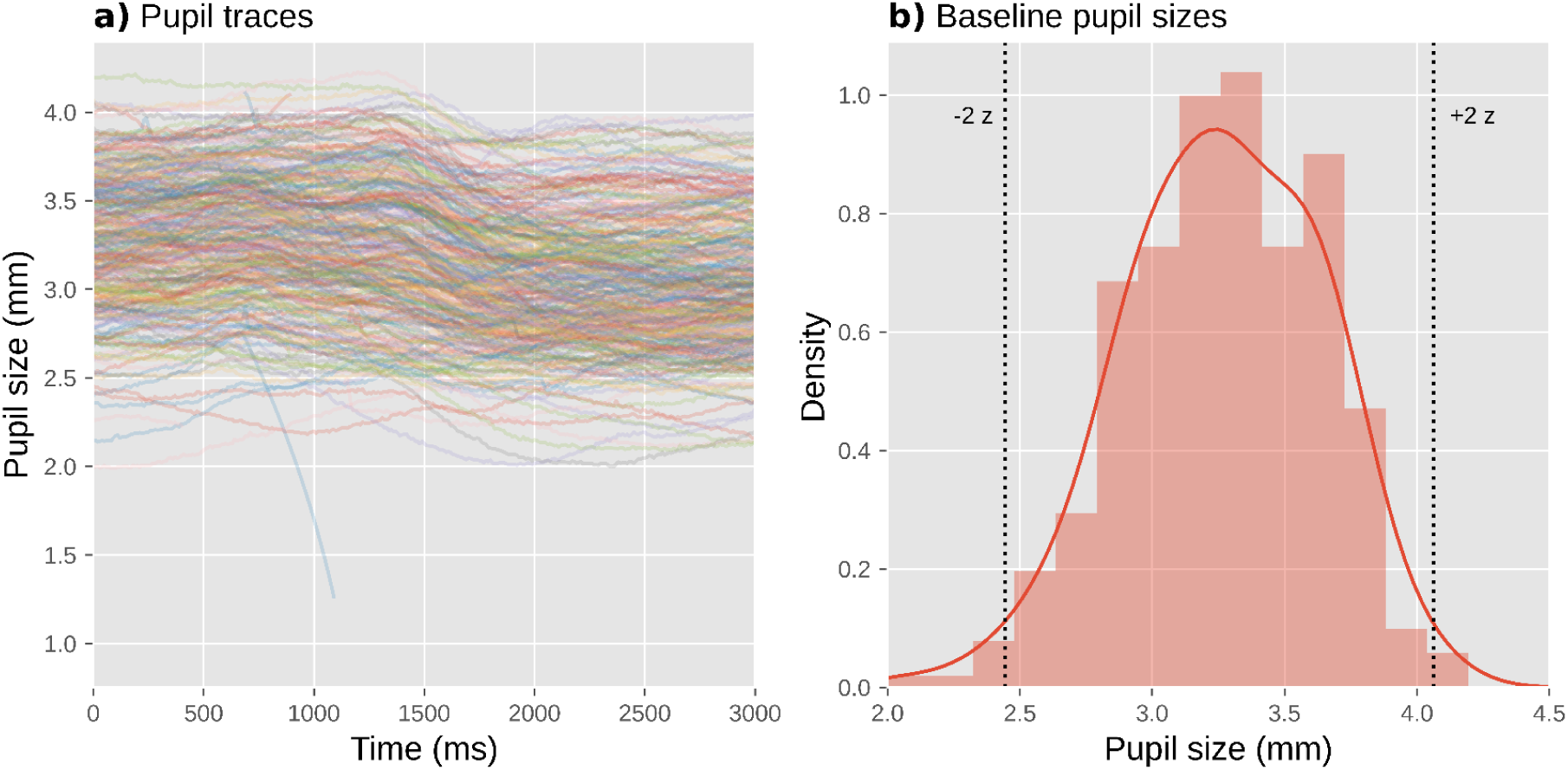
Visualization of data quality for a single participant from our example experiment. a) A plot with pupil size during all trials plotted as semi-transparent lines allows you to check visually for distortions, such as downward spikes. b) A histogram of baseline pupil sizes allows you to check visually for clusters of outliers.

A key sign of poor data quality is the presence of spikes that shoot downwards (but rarely upwards) from this tangle of lines: these spikes correspond to blinks or other recording artifacts that were not successfully interpolated or removed. A handful of such spikes can occur even in high-quality data (such as the downwards spike around 1000 ms in Fig. 7a), and this is not necessarily a cause for concern; however, if there are too many spikes, this adds substantial variability to the data, thus decreasing statistical power. What constitutes ‘too many’ is largely a matter of experience and tolerance, but as a rule of thumb no more than 5% of trials should show such spikes. If there are more spikes, then it is worthwhile to reconsider the procedure for interpolating and removing invalid data (see Interpolating or removing invalid data); for example, you could try out different parameters for the blink-reconstruction function to see if the number of spikes is reduced with certain parameter combinations. Unfortunately, the quality of blink reconstruction strongly depends on choosing the right parameters, and whereas the default parameters of DataMatrix `blinkreconstruct()` work well on 1000 Hz data as recorded by the EyeLink 1000 eye tracker, different eye trackers may require manual adjustment through trial and error.

Another sign of poor data quality is the presence of lines that are far above (but rarely below) the others, or that start from zero but then quickly (< 200 ms) shoot upwards: these lines correspond to trials on which there were artifacts during the baseline period, resulting in extremely small baseline pupil sizes (which should also be evident in the histogram of baseline pupil sizes) and thus extremely large baseline-corrected pupil sizes (Mathôt et al., 2018). Such trials add substantial variability to the data, thus decreasing statistical power. Fortunately, it is straightforward to identify and exclude such trials (see Trial exclusion based on baseline pupil size).

A third useful visualization is to plot the average number of blinks per trial, separately for each experimental condition and participant (Fig. 8). Participants differ considerably in how often they blink; in our example experiment, some participants rarely blink at all, while others blink several times on each trial. This is not in itself a problem; however, blink rate should not systematically differ between experimental conditions. If it does, then this is inherently problematic when interpreting differences in pupil size between experimental conditions, because pupil size is affected by blinks (Yoo et al., 2021). If you find differences in blink rate, then you may want to reconsider the experimental design in order to avoid this. If changing the design is not possible, then differences in blink rate should at least be transparently reported, such that readers and reviewers can assess for themselves whether these differences are likely to confound the results.

**Figure 8.**
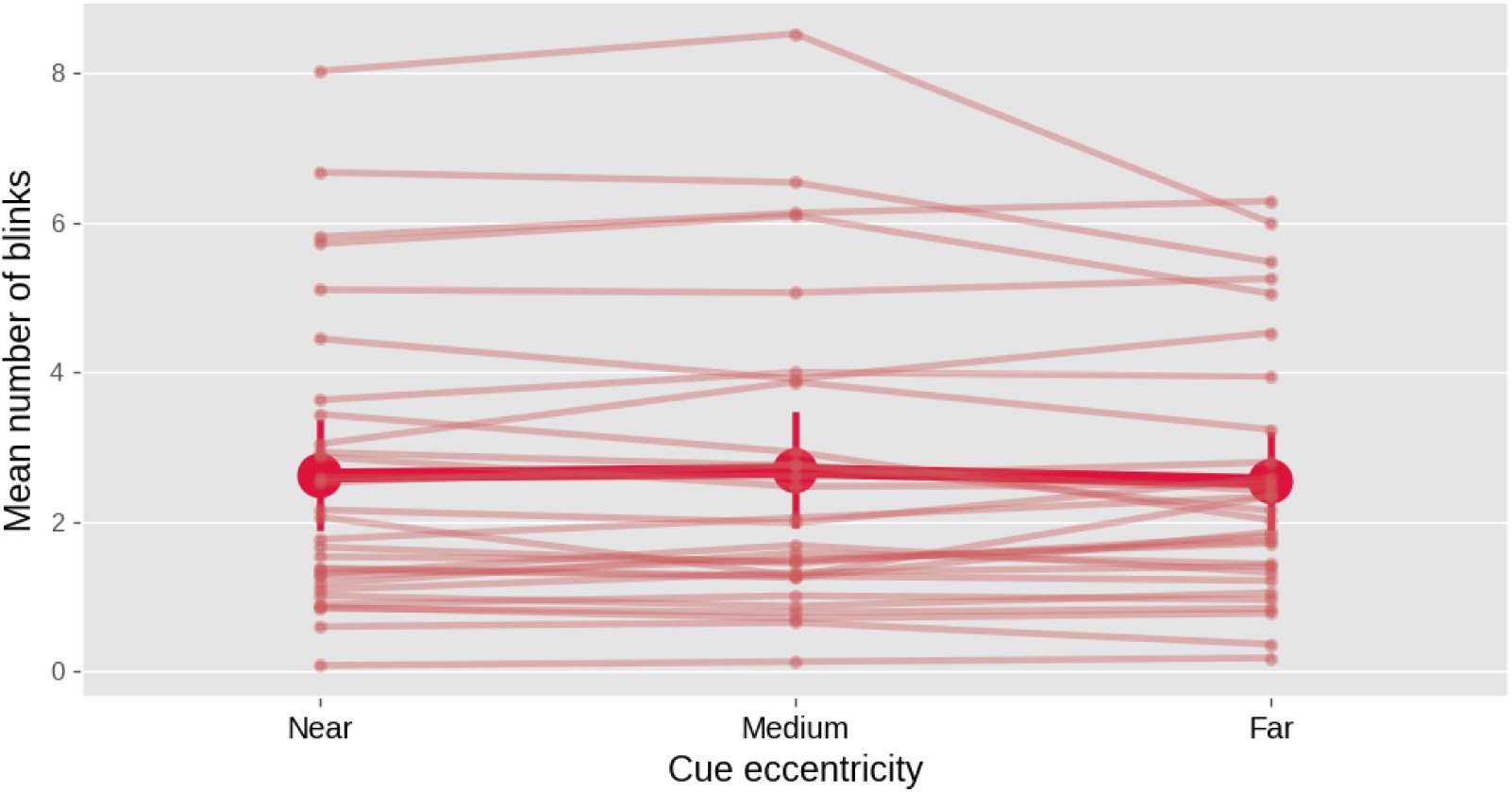
A plot with the mean number of blinks per trial as a function of experimental condition and participant allows you to check visually for systematic differences in blink rate between conditions. Error bars indicate 95% between-subject bootstrapped confidence intervals. Based on complete data from our example experiment.

Some eye trackers, such as the EyeLink that we used for our example experiment, automatically detect blinks during recording, in which case it is easy to create a plot like Fig. 8. For eye trackers that do not automatically detect blinks, you can use a custom blink-detection algorithm (e.g. Hershman et al., 2018). Alternatively, you can count the number of missing data points on each trial as an easy proxy for the number and duration of blinks (although of course missing data can also result from other factors, such as recording errors).

In our example experiment, there was no notable difference in blink rate between experimental conditions (Fig. 8), although there were substantial differences in blink rate between participants.

A fourth and final useful visualization is to show gaze position over time. In our example experiment, participants were instructed to keep their eyes fixated on a central fixation dot; therefore, the most relevant measure of eye position is a deviation from the display center in any direction. To visualize this, we plotted the absolute difference between the x coordinate and the horizontal display center, over time and averaged across trials but separately for each experimental condition, in one panel (Fig. 9a), and the absolute difference between the y coordinate and the vertical display center in another panel (Fig. 9b).

**Figure 9.**
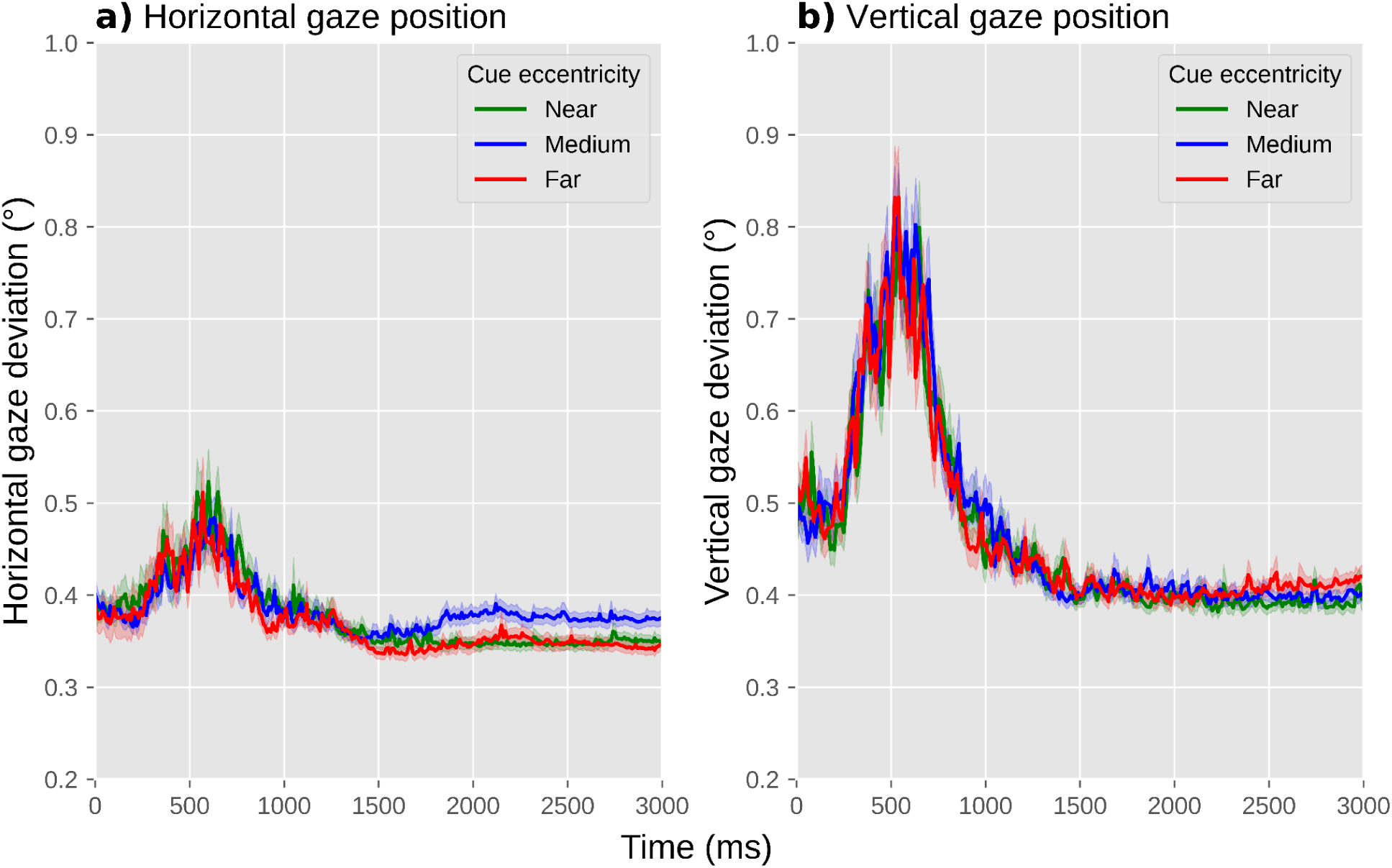
A plot with the deviation of horizontal (a) and vertical (b) eye position from the display center as a function of experimental condition allows you to visually check whether there are systematic differences in gaze deviation between conditions. Error shadings indicate grand standard error. Based on complete data from our example experiment.

When assessing differences in gaze position between experimental conditions, the situation is much the same as for blink rate: ideally, there should not be any systematic differences between experimental conditions in either horizontal or vertical gaze position; if there are, this may require a change to the experimental design, and if this is not possible, then these differences should be reported.

In our example experiment, there was no notable difference between conditions in absolute gaze deviation from the center (Fig. 9). Gaze deviation briefly increased around 500 ms after the onset of the cue, reaching a maximum deviation of about 0.5° for horizontal and 0.8° for vertical eye position, but again this occurred in all conditions. There was a hint of a slight increase in horizontal gaze deviation for the medium-cue-eccentricity condition relative to the other conditions, arising after about 1500 ms; this could have been a cause for concern if this difference in gaze deviation, however subtle, would have correlated with our predicted difference in pupil size, that is, if we had predicted pupil size to be smallest or largest in the medium-cue-eccentricity condition relative to the other conditions. However, this is not the case: we predict pupil size to increase as a function of increasing cue eccentricity, and therefore we can be fairly confident that, should we confirm our prediction, this result will not be confounded by gaze deviation.

### Excluding trials based on baseline pupil size

As described above, blinks and recording artifacts during the baseline period may result in very small baseline pupil sizes (see Baseline correction); in turn, this results in very large baseline-corrected pupil sizes, which add variability to the data and may substantially reduce statistical power (Mathôt et al., 2018). Therefore, trials with extreme baseline pupil sizes should be excluded from analysis.

To do so, you can first convert baseline pupil sizes to *z*-scores. This should be done separately for each participant, because what is an extremely small pupil size for one participant may be a fairly normal pupil size for another participant. Next, all trials where the *z*-scored baseline pupil size is larger than 2 or smaller than −2 are excluded (indicated by the vertical lines in Fig. 7b). This simple, predetermined criterion tends to effectively remove problematic trials. In the data from our example experiment, we excluded 402 trials (5.58%) based on this criterion.

### If necessary: excluding participants based on data quality

Data quality can differ substantially between participants, for example because of contact lenses, glasses, eye make-up, or other factors that reduce the ability of the eye tracker to record the pupil. In rare cases, this may be a reason to exclude a participant’s data from analysis altogether. Ideally, exclusion criteria should be specified in advance; however, because of the many hard-to-predict ways in which the quality of pupil-size data can be poor, you may still encounter participants that warrant exclusion for reasons that you did not foresee. Therefore, it is crucial to transparently report how many participants were excluded and why.

Fortunately, a few precautionary steps often prevent having to exclude any participants. First, make sure to properly set up the participant before the experiment. For example, make sure that the participant is well-positioned in front of the eye tracker, that the chin rest is comfortable, that the focus of the eye-tracking camera is adjusted (if applicable), that the eye tracker is well-calibrated, etc.

Second, decide in advance what to do with an experimental session if you suspect that the data will be unusable, for example because the eye tracker regularly fails to track the eye. One possibility is to abort such sessions altogether, so that you prevent yourself from having to decide post-hoc—and thus biased by whether you like the results—whether to include the data or not. (Assuming, of course, that incomplete data is never included, which is a general rule that most researchers, including ourselves, follow unless there is a good reason to do otherwise.) When aborting a session, consider that participants may take this as a sign of failure on their part, and it is therefore courteous to explain to them that eye tracking is prone to technical issues even if participants perform well. Another possibility is to make a note in a laboratory log book that eye-movement and pupil data for particular participants should be discarded; this has the advantage that other data, such as behavioral responses, can still be analyzed. However, the decision to include or exclude a participant’s data, which is often a subjective decision, should ideally be made before the data have been analyzed further, again in order to avoid this decision from being biased by whether or not you like the participant’s results.

Third, use a statistical technique that is able to deal with large differences in observations between participants; that way, you do not have to exclude participants if a substantial number of trials is excluded due to, for example, extremely small baseline values (see Excluding trials based on baseline pupil size).

In our example experiment, we did not exclude any participants from analysis.

## Visualization and statistical analysis

From a statistical perspective, most cognitive-pupillometry experiments are interested in the following question: do any of my experimental manipulations affect pupil size at some moment during an interval of interest? This is also the case for our example experiment, in which we were interested in whether cue eccentricity affects pupil size at some moment between the presentation of the cue and the target.

More complex measures can also be derived from pupil-size data. For example, Reilly et al. (2021) discuss how a pupil-dilation response can be quantified as a time-to-peak (the time it takes for the pupil to reach its maximum size), base-to-peak (the maximum size that the pupil reaches), and several other measures; Fink et al. (2021) discuss how the synchronicity between pupil size and external events, such as rhythmic beats, can be quantified; and Wierda et al. (2012) introduced a ‘deconvolution’ technique to quantify how the (sluggish) pupil response is affected by individual events that are part of a rapid sequence of events, such as stimuli in a rapid-serial-visual-presentation (RSVP) stream. All of these approaches warrant a full discussion in their own right, and we therefore refer to these respective articles for further reading. Here we will limit ourselves to the modest, yet often very useful, question of: does my manipulation affect pupil size at all?

### Multiple comparisons in analysis of pupil-size data

For our purpose, the main factor that complicates analysis of pupil-size data, as compared to response-time and most other behavioral data, is that pupil size is a continuous signal or ‘time series’; that is, the data does not contain a single dependent variable, but rather a series of dependent variables, one for each time sample, that reflect how pupil size changes over time.

In our example experiment, there were 300 such values, one for each 10 ms of our 3000 ms cue-target interval. This means that we could, in principle, conduct 300 statistical tests, one for each sample, and then see if one of these tests is ‘significant’. Specifically, we could test 300 separate linear mixed effects models with (baseline-corrected) pupil size for one specific 10 ms sample as dependent variable, cue eccentricity as fixed effect, and participant as random effect. Doing so reveals *p* < .05 for the effect of cue eccentricity at 125 samples.

However, this approach clearly raises the issue of multiple comparisons: when conducting so many statistical tests, there is a high chance that some of these tests will give significant but bogus effects; using statistical terminology, there will be a high chance of *spurious effects* or *false alarms* or *type I errors*, or the *family-wise error rate* exceeds the intended alpha level. To complicate matters further, pupil-size data is “auto-correlated”: it changes very little from one 10 ms sample to the next. Because of this auto-correlation, you should not correct *p* values with traditional techniques that assume that observations are uncorrelated, such as Bonferroni correction. To mitigate the issue of multiple comparisons, some researchers, including ourselves, have used an additional criterion, such as that *p* < .05 should hold for at least 200 contiguous milliseconds (e.g. Mathôt et al., 2017; see also Hershman et al., 2022 for a similar approach using Bayes Factors). This improves the issue somewhat, but it is not a formal way to correct for multiple comparisons; therefore, we no longer recommend this.

Sophisticated techniques for analyzing auto-correlated time series have been developed for functional magnetic resonance imaging (fMRI), electroencephalography (EEG), and magnetoencephalography (MEG; Bowman et al., 2020; Bullmore et al., 1999; Kriegeskorte et al., 2009; Maris, 2004; Maris & Oostenveld, 2007); and although the specifics differ, the general principles that have been developed in these fields also apply to pupil-size data. Here we will focus on three techniques: using a predetermined time window, cluster-based permutation testing, and—our preferred technique—cross-validation.

### Using a predetermined time window

If you have a strong prediction about when an effect should arise, you can conduct a single statistical test on the mean pupil size during the interval during which you predict the effect to emerge.

For our example experiment, we could derive a prediction from previous experiments that found that a voluntary shift of covert visual attention towards a bright or dark surface affects pupil size from about 750 ms after cue onset until the end of the trial (Mathôt et al., 2013). Based on this, we might predict that the effect of attentional breadth on pupil size similarly arises from 750 ms after the onset of the cue, and persists until the end of trial. We could then determine the mean (baseline-corrected) pupil size during this interval for each trial, and conduct a linear mixed effects analysis with Mean Pupil Size as dependent variable, Cue Eccentricity (−1: near, 0: medium, 1: far) as fixed effect, and Participant as random effect. Using an alpha level of .05, this test shows an effect of Cue Eccentricity (*z* = 2.28, *p* = .023) such that—as predicted—pupil size increases with increasing attentional breadth. A direct visualization of this test would be a line or bar plot that displays mean baseline-corrected pupil size as a function of cue eccentricity, with error bars that show the standard error across all trials (Fig. 10). However, even when analyzing predetermined time windows, it is still informative to also include a figure that shows how pupil size changes over time (Fig. 12), because this often provides important information that is lost in line or bar plots.

**Figure 10.**
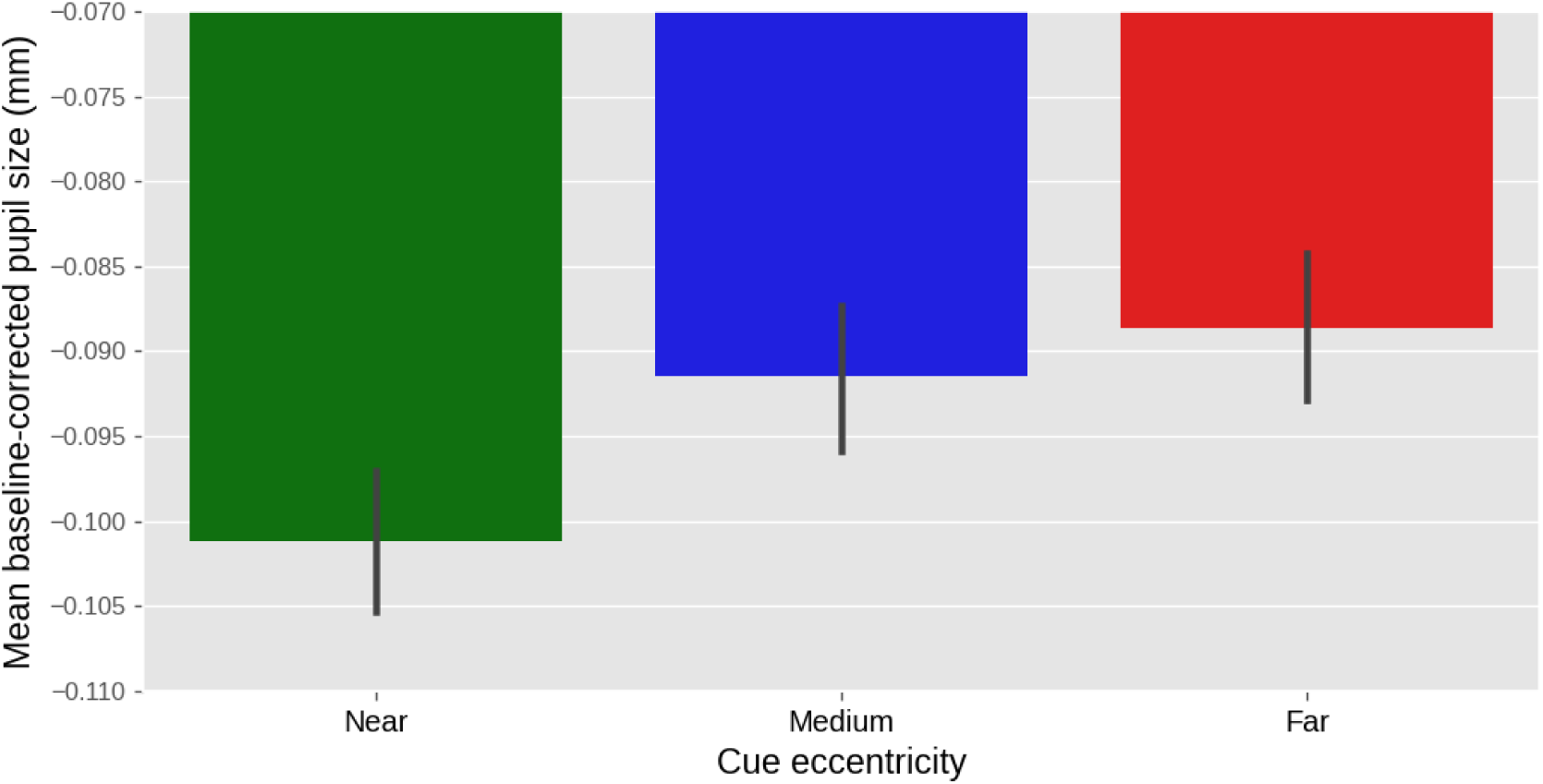
Results from our example experiment. Mean baseline-corrected pupil size during the 750 ms - 3000 ms window after cue onset as a function of cue eccentricity. Values are negative due to an overall pupil constriction relative to the baseline period (see also Fig. 12). Error bars indicate grand standard error.

The advantage of analyzing a predetermined time window is that there is no need to compensate for multiple comparisons at all. The disadvantage is that, unless you are conducting a direct replication of a previous experiment, it is difficult to know in advance exactly at which point in time an effect, if it exists, will emerge: this depends on inter-individual variability, the stimulus display, task demands, and other factors (see also Bowman et al., 2020 for a similar point for EEG experiments).

### Cluster-based permutation testing

Let’s go back to the 300 separate tests that we conducted on the data from our example experiment. In this data, we found a cluster of 125 consecutive samples for which *p* < .05 (i.e. all 125 samples for which *p* < .05 comprised a single cluster). The crucial question is then, given that we are likely to find *some* samples for which *p* < .05, how likely are we to find a cluster of the size that we observed or larger, assuming that the null hypothesis is true? Phrased differently, what is the *p* value of this cluster?

A cluster-based permutation test answers this question by randomly shuffling the condition labels on a trial basis and then running the same analysis again (Bullmore et al., 1999; Maris & Oostenveld, 2007). Any cluster of samples for which *p* < .05 is then by necessity a false alarm. This procedure of shuffling and analyzing is repeated a large number of times (e.g. 1000 times), which results in a distribution of false-alarm clusters. The *p* value for the actual cluster (i.e. the cluster size based on the unshuffled data) is then the proportion of false-alarm cluster sizes that are larger than or equal to the actual cluster size.

Cluster-based permutation tests are elegant and effective. However, they are not feasible when using linear mixed effects models to determine *p* values for individual samples, because the time it would take to conduct the required number of tests is prohibitive. A single linear mixed effects model takes at least two seconds to run. To conduct a 1000-fold cluster-based permutation test for a time series of 300 samples would take at least 1000 (permutations) × 300 (samples) × 2 (seconds) = 600,000 seconds, or one week (!) to compute.

The computational time required for a cluster-based permutation test can be vastly reduced by using a different underlying statistical technique, such as a repeated measures ANOVA, to determine *p* values for individual samples. However, this would also result in decreased statistical power due to the fact that a repeated measures ANOVA is conducted on aggregated data (i.e. mean pupil size per participant and condition), whereas a linear mixed effects analysis is conducted on individual trials (Brysbaert & Stevens, 2018). For this reason, we prefer to preserve the statistical power of linear mixed effects models, and combine this with cross-validation to control for multiple comparisons.

### Cross-validation testing

Let’s again go back to the 300 separate tests that we conducted on the data from our example experiment. Now that we have seen the data, we have a better idea of where to look: the sample for which the *p* value was lowest (sample 261, *p* = .002). However, if we would actually do this, we would perform a circular analysis, in the sense that we would use the same data to both identify and test our result, which results in a very high chance of a false alarm (Bowman et al., 2020; Kriegeskorte et al., 2009).

Cross-validation is a general approach to avoid circularity in analyses by using one part of the data (the training set) to localize an effect, and another part of the data (the test set) to test the effect at the location that was identified in the training set. The Python library `time_series_test`^3^ implements cross-validation with linear mixed effects modeling in a way that is suitable for pupil-size data, and the general procedure is visualized in Fig. 11. Specifically, 75% of the data is used for the training set, and the remaining 25% of the data is used for the test set (using four-fold cross-validation, which is the default); the data is (by default) split in an interleaved fashion, such that three subsequent rows go into the training set, then the next row goes into the test set, the next three rows go into the training set, and so forth. Interleaved splitting is done across the entire dataset, without taking into account how conditions are distributed across trials. A linear mixed effects model is then conducted for each sample of the training set. The sample that yields the highest *z* value in the training set is used as the sample-to-be-tested for the test set. This procedure is repeated four times, using a different training set each time, until all samples have been part of a test set, and a sample-to-be-tested has therefore been determined for the entire dataset. Finally, a single linear mixed effects model is conducted using the sample-to-be-tested for each trial as a dependent measure. This means that the dependent variable consists of a column of (baseline-corrected) pupil-size values that correspond to different samples for different trials.

**Figure 11.**
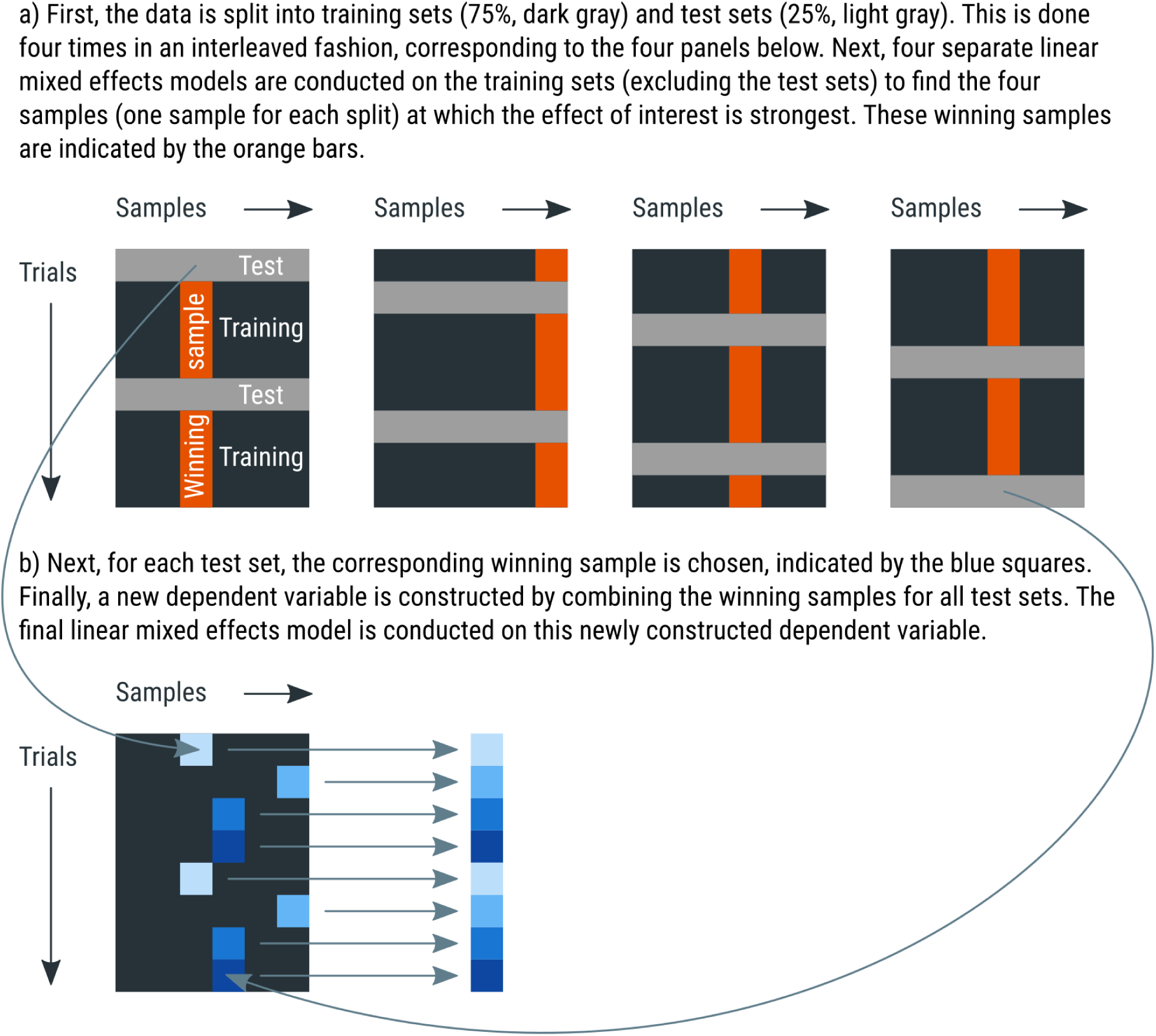
A schematic explanation of four-fold cross-validation. See main text for details.

For models with multiple fixed effects, the cross-validation procedure is repeated for each main effect and interaction in the model. This is necessary because different effects may arise at different moments.

Using an alpha level of .05, we ran a cross-validation analysis on our predetermined time-window of 750-3000 ms, and found an effect of Cue Eccentricity (*z* = 2.92, *p* = .004, tested at samples 261 and 297) such that—as predicted—pupil size increases with increasing attentional breadth. An appropriate visualization of this test could be a ‘trace plot’ that shows mean pupil size as a function of time with cue eccentricities as separate lines (Fig. 12). The mean of the to-be-tested samples can then be shown as a vertical marker indicating when, approximately, the effect of cue eccentricity occurs most strongly, in this case at sample 279.

**Figure 12.**
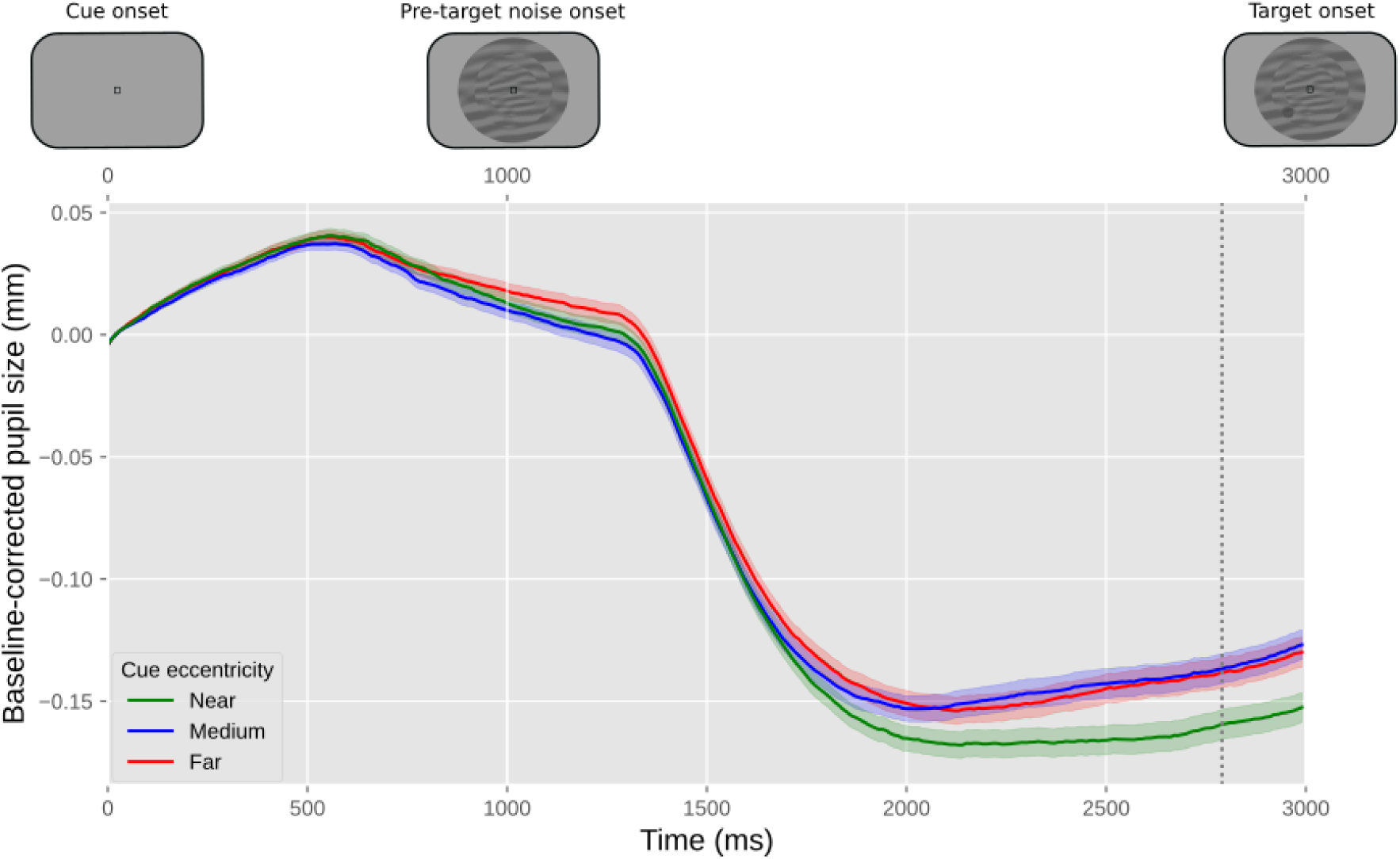
Results from our example experiment. The y-axis represents baseline-corrected pupil size. The lower x-axis represents time in milliseconds since cue onset. The upper x-axis represents the order of the events in the experiment. Differently colored lines represent the three cue eccentricities (far, medium, near). Error bands indicate grand standard error. The vertical line indicates the mean of the samples at which the effect of cue eccentricity was tested.

When visualizing the data like this, it also becomes apparent that the onset of the noise stream is followed by a pronounced pupil constriction. This constriction has a latency of around 250 ms, which is characteristic for constrictions induced by visual stimulation. This effect is not of primary interest to us, but is far larger than the effect of our experimental manipulations, which highlights the importance of keeping visual stimuli constant between conditions (see Stimuli should ideally be constant between conditions).

The main advantage of cross-validation, as compared to cluster-based permutation testing, is that it requires far fewer mixed effects models to run. For our example study, and again assuming at least two seconds of computational time for each linear mixed effects model, the test requires at least 4 (splits) × 300 (samples) × 2 (seconds) = 2,400 seconds, or 40 minutes to compute. The test can be sped up considerably by using a so-called random-intercept only model for the localization phase (which is the default), and include by-participant random slopes, which is recommended by many statisticians (Barr et al., 2013), only for the final model that gives the final *z* and *p* values. For our example data and our test system, this reduces the computational time to about 5 minutes. Another advantage of cross-validation is that the outcome is deterministic—unlike the outcome of cluster-based permutation testing—at least when using a deterministic ‘interleaved’ method of splitting the data into test and training sets (which the default); that is, you always get the exact same outcome for the same data and analysis parameters.

## Conclusion

Here we have provided a comprehensive, hands-on guide to cognitive pupillometry, which is the general approach of using pupil size as a measure for various cognitive processes. The guidelines are summarized in Fig 1. We have focused on (to us) ‘typical’ cognitive-pupillometry experiments, which are trial-based experiments in which a task-evoked pupil response to a stimulus is the measure of interest. We have outlined six basic principles for designing such experiments (see Experimental design):

- Stimuli should ideally be constant between conditions (“the Hillyard principle”)
- Eye position should ideally be constant between conditions
- Trials should ideally be slow-paced (have stimuli of interest be followed by 2 - 3 s of recording period, and use an intertrial interval of at least 3 s)
- Pupil size should ideally be measured while participants do nothing
- Ambient lighting should ideally be intermediate (± 30 lux) and matched to display brightness (background luminance around 30 cd/m^2^)
- All data should ideally be stored in a single data file per participant

We further provided crucial steps for preprocessing pupil-size data, where ‘preprocessing’ refers to all steps involved in transforming raw data, as recorded during the experiment, into a format that is suitable for statistical analysis and visualization (see Preprocessing). We have also provided example Python code that illustrates these steps. In order (and the order matters!), the preprocessing steps are:

- Parsing raw data into an analysis-friendly data structure
- Interpolating or removing missing and invalid data (e.g. due to blinks)
- If necessary: downsampling
- If necessary: converting pupil size from arbitrary units to millimeters of diameter
- Baseline correction
- Verifying and visualizing data quality
- Excluding trials based on baseline pupil size
- If necessary: excluding participants based on data quality

Finally, we have discussed how to conduct statistical tests on pupil-size data (see Visualization and statistical analysis), highlighting the issue of multiple comparisons, which arises because pupil size consists of multiple observations over time, and it is often not known in advance at which time point an effect is expected to emerge. We have suggested three appropriate statistical approaches: conducting a single test on a predetermined time window; conducting a cluster-based permutation test; and conducting a cross-validation test. We have suggested that cross-validation is in many cases the preferred approach, because it does not require a time window to be determined a priori, and (unlike a cluster-based permutation test) it can be combined with linear mixed effects modeling without becoming prohibitively computationally intensive.

Finally, we have emphasized that our guidelines are illustrative rather than prescriptive: they are intended to provide researchers with a solid starting point for conducting cognitive-pupillometry experiments.

## Open practices statement

Experimental data and analysis scripts for this methods article are publicly accessible at https://osf.io/659pm/.

## Footnotes

1 https://github.com/smathot/python-eyelinkparser

2 https://pydatamatrix.eu/

3 https://github.com/smathot/time_series_test

